# Conformational ensemble dependent lipid recognition and segregation by prenylated intrinsically disordered regions in small GTPases

**DOI:** 10.1101/2023.08.11.553039

**Authors:** Mussie K. Araya, Alemayehu A. Gorfe

## Abstract

We studied diverse prenylated intrinsically disordered regions (PIDRs) of Ras and Rho family small GTPases using long timescale atomistic molecular dynamics simulations in an asymmetric model membrane of phosphatidylcholine (PC) and phosphatidylserine (PS) lipids. We show that conformational plasticity is a key determinant of lipid sorting by polybasic PIDRs and provide evidence for lipid sorting based on both headgroup and acyl chain structures. We further show that conformational ensemble-based lipid recognition is generalizable to all polybasic PIDRs, and that the sequence outside the polybasic domain (PBD) modulates the conformational plasticity, bilayer adsorption, and interactions of PIDRs with membrane lipids. Specifically, we found that palmitoylation, the ratio of basic to acidic residues, and the hydrophobic content of the sequence outside the PBD significantly impact the diversity of conformational substates and hence the extent of conformation-dependent lipid interactions. We thus propose that the PBD is required but not sufficient for the full realization of lipid sorting by prenylated PBD-containing membrane anchors, and that the membrane anchor is not only responsible for high affinity membrane binding but also directs the protein to the right target membrane where it participates in lipid sorting.

## Introduction

Cells use primarily two types of signals to target proteins to membrane surfaces: modular protein domains and lipid-based motifs^1^. Modular membrane targeting protein motifs such as C1, C2, PH, and BAR domains have been studied extensively (ref.^1,2^ and references therein). Another class of a well-studied membrane targeting motif utilizes a combination of an amphipathic helix and a cluster of basic residues for lipid binding (e.g.,^3,4^). Many proteins also use intrinsically disordered regions (IDRs) for membrane binding, and rules are emerging regarding their lipid recognition propensities such as preferences of basic amino acids for anion membranes^5^. While most lipid-based membrane targeting motifs also harbor IDRs^2,6,7^, insights derived from non-lipidated and autonomously membrane interacting IDRs are not directly applicable to cases where co- or post-translational lipid modification is an absolute requirement for membrane binding. This is the case, for example, in the Ras superfamily of small GTPases. As a result, our understanding of lipid recognition by lipidated IDRs remains limited despite useful insights from molecular dynamics (MD) simulation studies of Ras proteins^8–10^. An important open question of particular interest to the field is how a lipidated IDR might engages and sorts distinct membrane lipid species.

Membrane binding and therefore cellular function of most Ras superfamily proteins and many other intracellular signaling proteins requires post-translational prenyl (farnesyl or geranylgeranyl) lipid modification at a C-terminal cysteine residue^10–12^. High affinity membrane binding is often achieved by complementing the prenyl modification by a proximal polybasic domain (PBD) or an additional lipid modification^13–15^. The PBD is located within a ∼10-20 residue-long flexible region, which we refer to as the <u>p</u>renylated intrinsically <u>d</u>isordered region (PIDR). It is comprised of 4-8 contiguous or semicontiguous (i.e., separated by no more than two amino acids) lysine and/or arginine. Typical examples of PBD-containing PIDRs in the Ras superfamily include the plasma membrane (PM) associated KRAS4B (hereafter KRAS) and RhoA whose mutation or overexpression is known to cause cancers^10–12,16^. These proteins achieve high affinity membrane binding through a combination of van der Waals (vdW) interactions of the prenyl chain with membrane lipid acyl chains and electrostatic interactions of the PBD with the headgroup of anionic lipids, such as phosphatidylserine (PS) in the inner leaflet of the PM^13,14,17^. In addition, the membrane anchor plays a crucial role in the spatial distribution of RAS proteins on the PM^11,13,15,18–20^. Intriguingly, the specific sequence position of the basic residues in the PBD and the identity of the prenyl group were found to be important determinants of membrane binding and lipid sorting by KRAS^21–25^ and Rac1^26^. Moreover, we recently showed that the interactions of the KRAS PBD with PS lipids varies with its backbone conformational dynamics^22,27^, and that charge-preserving mutations within the PBD caused changes in lipid recognition^27^. We therefore proposed that the conformational plasticity of the PIDR allows for the PDB and the prenyl group to act as a combinatorial code for lipid sorting by all PBD-containing prenylated small GTPase^27^. A primary goal of the current work is to formally test this hypothesis.

PBD-containing PIDRs differ in sequence length and net positive charge as well as in the ratio of Lys to Arg residues within the PBD. Differences within the region N-terminus to the PBD include the presence or absence of additional lipid modifications, hydrophobic content, and the ratio of basic to anionic amino acids. Another factor to consider is the spacing of the PBD from the prenylation site. If lipid sorting is sensitive to structural fluctuations of the PIDR as found in KRAS^25^, how would these differences affect the membrane localization and lipid interaction of PBD-containing PIDRs? A second goal of this study is to examine the impact of these variations on PIDR structure, dynamics, and membrane binding. To this end, we studied five diverse PM targeting, PBD-containing PIDRs in a compositionally asymmetric PC/PS bilayer. These include the 19-residue long PIDRs of Rap1A and Rap1B from the Ras family and Rac1 and Cdc42b from the Rho family. These PIDRs differ in sequence within the PBD and in the spacing of the PBD from the prenylation site, as well as in additional lipidation, hydrophobic residue content, and ratio of basic to acidic residues outside the PBD. We have also examined shorter PIDRs, one from RhoA with a similar PBD to those of the other PIDRs and another from Rheb that does not have a PBD.

These systems were investigated using atomically detailed MD simulations of 58μs aggregate effective time. The results confirm the key role of backbone conformational plasticity in lipid recognition, and further show that the sequence outside the PBD plays an important role in the adsorption of the entire peptide onto the host anionic monolayer, lipid sorting, and interaction patterns. Moreover, while the spacing of the PBD from the prenyl group has a negligible effect, palmitoylation as well as the ratio of basic to acidic residues and the hydrophobic content of the sequence N-terminal to the PBD significantly impact the diversity of conformational substates and hence the extent of conformation-dependent PS interactions.

## Methods

### Model selection

Our goal in this work was to test the hypothesis that backbone conformational plasticity of PBD-containing PIDRs is correlated with their lipid recognition profile, and that this ensemble-based lipid recognition is tuned by the sequence length and composition of the PIDR as well as by the charge and spacing of the PBD from the prenylated cysteine. To test this hypothesis, we selected five diverse PBD-containing PIDRs as model systems, representing the membrane targeting motifs of RhoA, Rap1A, Rap1B, Rac1 and Cdc42b (Table 1). The C-terminal Cys of all five proteins is post-translationally modified by geranylgeranylation (GG) and carboxymethylation (Fig 1A; Table S1). The PIDRs of Rap1A, Rap1B, Rac1 and Cdc42b have the same sequence length (SL = 19) but differ in the net charge (Np = 6, 4, 6, 4) and Lys to Arg ratio (K:R = 6:0, 3:1, 4:2, 2:2) within the PBD (Table 1 and Fig 1A; see also Table S1). They also differ in the region N-terminus to the PBD, including in terms of hydrophobic content defined as the ratio of hydrophobic to total number of residues (Nh = 0.33, 0.45, 0.83, 0.70), the ratio of basic to acidic amino acids (B:A = 3:1, 3:0, 1:1, 0:4), and the presence of a palmitoylated cysteine in Rac1. The RhoA PIDR is shorter (SL = 10) and lacks a hydrophobic N-terminal extension, but it has a comparable PBD to Rap1B (Np = 5 vs. 4; K:R = 3:2 vs. 3:1). These five proteins also differ in the spacing of the PBD from the prenylation site, *S*, defined as the number of residues between the last residue of the PBD and the prenylated cysteine. For control, we included the farnesylated (Farn) membrane targeting motif of Rheb that has a similar SL to RhoA but lacks a PB sequence (Table 1 and Fig 1A).

**Figure 1.**
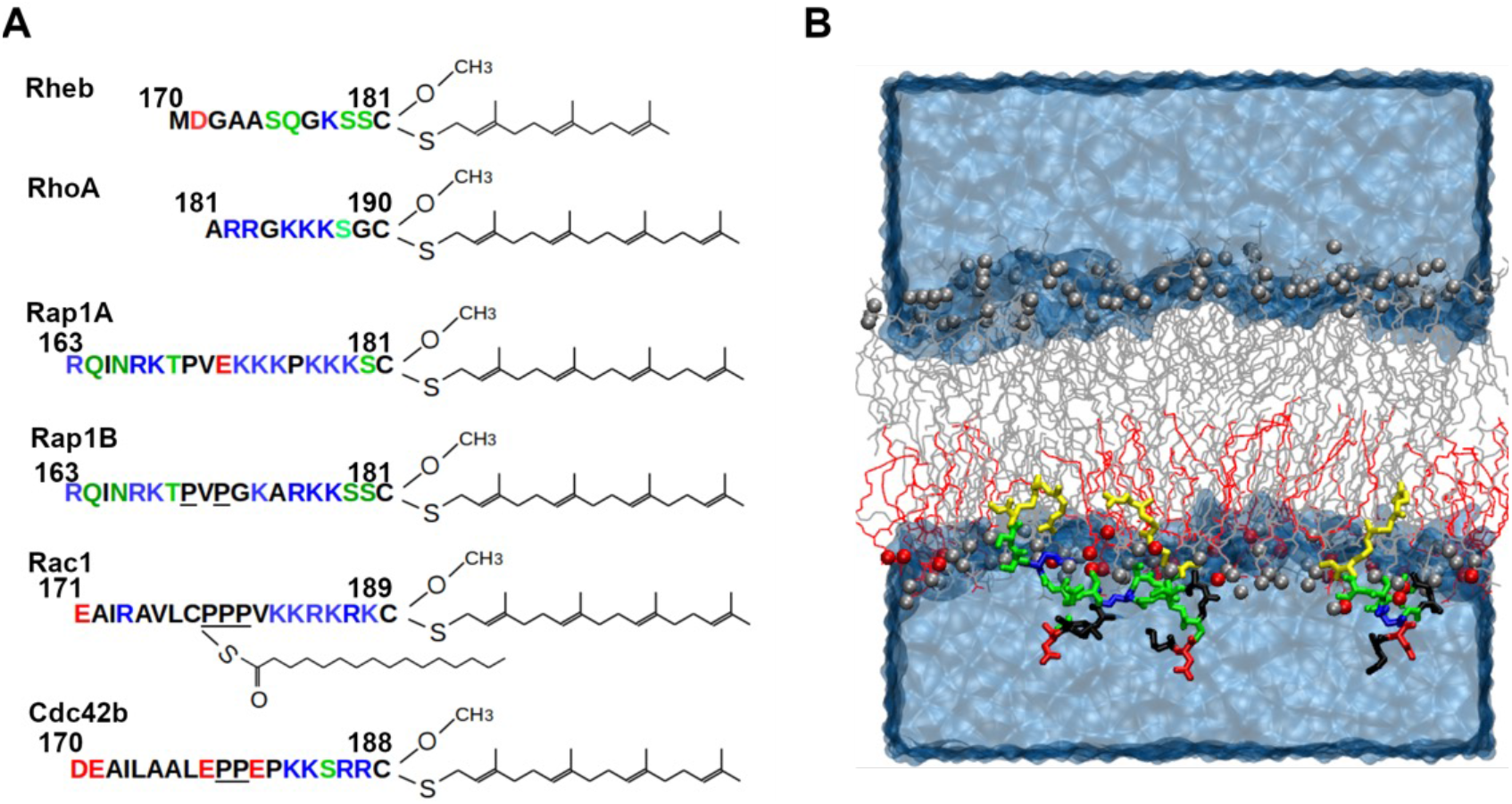
Prenylated intrinsically disordered regions of small GTPases investigated in this work and simulation setup. (A) The amino acid sequence and lipid-modification of the simulated PIDRs (color code: basic blue; acidic red; polar green; hydrophobic black). The C-terminal Cys residue is carboxymethylated following farnesylation (Rheb) or geranylgeranylation (all others). Rac1 is also palmitoylated at residue 178. 2-3 sequentially proximal prolines are underlined. (B) An example of the simulation setup, with three peptides embedded in the mixed-lipid leaflet of an asymmetric model membrane composed of POPC (grey) and POPS lipids (red). Lipid phosphorus atoms are shown in vdW spheres and the peptides (in this case Rheb) in licorice colored as in panel A, except for the prenylated Cys residues which are in yellow. Water and ions are shown as a continues blue surface.

**Table 1:** The membrane-targeting prenylated intrinsically disordered regions (PIDRs) of small GTPases investigated in this work.

| PIDR | Sequence length (SL) | Lipid modification | | PBD | | N-term to PBD | | C-term to PBD | # of atoms | Sim. Length ($\mu s \times pept^*$) |
| --- | --- | --- | --- | --- | --- | --- | --- | --- | --- | --- |
|  |  | *Prenyl* | *Palm* | *Np* | *K:R* | *Nh* | *B:A* |  |  |  |
| **Rheb** | 12 | Farn | - | - | - | - | - | - | 66459 | 3 x 3 |
| **RhoA** | 10 | GG | - | 5 | 3:2 | - | - | 2 | 63708 | 3 x 3 |
| **Rap1A** | 19 | GG | - | 6 | 6:0 | 0.30 | 3:1 | 1 | 61000 | 10 x 3 |
| **Rap1B** | 19 | GG | - | 4 | 3:1 | 0.45 | 3:0 | 2 | 64063 | 10 x 3 |
| **Rac1** | 19 | GG | yes | 6 | 4:2 | 0.83 | 1:1 | 0 | 72887 | 10 x 1 |
| **Cdc42b** | 19 | GG | - | 4 | 2:2 | 0.70 | 0:4 | 0 | 63079 | 10 x 3 |

### Modeling PIDR initial structures

No experimental structure is available for any of the PIDRs studied here. Therefore, we attempted to use SWISS-MODEL (https://swissmodel.expasy.org) and PEPFOLD (https://bioserv.rpbs.univ-paris-diderot.fr/services/PEP-FOLD/) servers to obtain consensus predictions of initial structures. However, only PEPFOLD yielded a complete structure for all peptides. Some of the predicted structures had a short helix but we selected models with the least amount of secondary structure with the assumption that the PIDRs are unstructured in solution. We then used CHARMM-GUI^28,29^ to attach a GG or Farn lipid and oxo-methylate the C-terminal cysteine. For Rac1, a palmitoyl lipid was also added to Cys178 (Fig 1A).

### PIDR-bilayer system construction

The inner leaflet of the PM where most of the PIDRs localize is enriched both in PS and phosphatidylethanolamine (PE) lipids. However, we showed recently that when compared with a PC-PC/PS bilayer, the presence of PE in an asymmetric PC-PC/PS/PE bilayer did not significantly alter the dynamics and membrane interactions of the KRAS PIDR^6^. Therefore, here we used the simpler binary bilayer in which PS lipids are distributed only to the monolayer where peptides are bound. An initial model of such a membrane was built as described recently^6^, with one leaflet containing 33 POPS and 77 POPC and the other 108 POPC lipids. This ensured area symmetry also considering the volume of the peptide lipids embedded in the mixed-lipid monolayer. To build initial PIDR-bilayer complexes, about 5 terminal carbon atoms of the prenyl chain were manually inserted into the hydrophobic core of the mixed-lipid leaflet. In this manner we attached three peptides per system to triple sampling with the computational cost of a single peptide (peptides were spaced 30-50Å center-of-mass distance apart to avoid self-interaction). The resulting PIDR-bilayer systems were solvated with TIP3P waters and neutralized by adding Na^+^. The final systems ranged in size between 61,000 and 72,887 atoms (Table 1). An example of the final setup is shown in Fig 1B.

### Molecular dynamics simulation and trajectory analysis

Each system was energy-minimized, and equilibrated as described previously^6^, and then subjected to a production run of 15-23ns using the NAMD program^30^ and the CHARMM36m force field^31^. The longer PIDR systems were then simulated for 10μs and the shorter ones for 3μs each on Anton 2 (Table 1), with trajectories written out every 100ps for analysis. Bilayer and protein structural and dynamic properties were analyzed as described in previous reports^6,8,32^. This included monitoring standard measures such as root mean square deviation (RMSD), radius of gyration (Rg), frequency of hydrogen bond (HB) and van der Waals (vdW) contacts, as well as bilayer thickness and area per lipid. Area per lipid (APL) was calculated from the area of the simulation box at the pure-POPC monolayer, and bilayer thickness using the average z-distance between the electron dense phosphorus (P) atoms at the two monolayers (P-P). Bilayer insertion depth, *I*, was calculated as the z-distance of the center-of-mass (COM) of the prenyl chain from the bilayer COM. HB was defined using donor-acceptor distance cutoff of 3.1Å and angle 30°, and vdW contact using a 4.0Å cutoff between pairs of carbon atoms. As in previous work^6,32^, we used multiple reaction coordinates (RCs) to characterize the highly dynamic PIDRs. These included global measures such as Rg, RMSD, and orientation relative to the membrane normal, as well as local features such as pseudo dihedral angles (Φ) involving virtual bonds between four consecutive C*α* atoms. Φ was computed for every four consecutive residues and the one that most clearly classified the simulated conformers into sub-ensembles was used for subsequent analyses. In Rac1, only one peptide has fully inserted into the bilayer and thus the total trajectory analyzed was 10µs. For the rest, three peptide trajectories were concatenated and the resulting 9µs (Rheb and RhoA) or 30µs (Rap1A, Rap1B and Cdc42b) effective time trajectories were used to compute equilibrium properties. We used R^33^ and VMD^34^ scripting for analysis.

## Results

We used long timescale atomistic MD simulations (aggregate effective time = 58µs) to investigate the membrane binding behavior, structure, and dynamics of prenylated intrinsically disordered regions (PIDRs) representing a diverse set of polybasic membrane targeting motifs of small GTPases (Table 1). These included Rheb, Rap1A and Rap1B from the Ras family and RhoA, Rac1 and Cdc42b from the Rho family. Each PIDR was simulated in an asymmetric PC-PC/PS bilayer to mimic the PM inner leaflet that is enriched with PS lipids.

### Convergence of the simulations

As in previous observations on a PC-PC/PS bilayer of the same lipid composition^6^, each of the current simulations equilibrated relatively quickly (within ∼0.5-2μs). This is exemplified in Fig 2A by the time evolutions of APL and P-P in the Rheb simulation, and Fig S1 shows a similar trend in all systems. Bilayer stabilization was fast in Rheb and RhoA and somewhat slow in Rap1A and Rac1 (Fig S1). This shows that the size and complexity of the PIDR affects the speed of bilayer equilibration. We therefore simulated the shorter and fast adsorbing Rheb and RhoA PIDRs for 3µs and the longer and hence harder to equilibrate Rap1A, Rap1B, Rac1 and Cdc42b for 10µs each (Table 1). As already noted, each simulation was started with the prenyl chain partially inserted into the bilayer. The plots of *I* versus simulation time (Figs 2B and S2) show that this setup allowed for a fast and complete insertion of the prenylated and oxo-methylated C-terminus, consistent with previous observations^6,32,35–39^. Similarly, the time evolution of backbone RMSD (Figs 2B and S2) show equilibration of the simulations and convergence of the peptide structures. Together with the snapshots in Fig 3, these results demonstrate that bilayer adsorption and structural reorganization occurred within 0.5-1µs in all except the palmitoylated Rac1, which took longer (2.2µs). We therefore excluded the first 0.5µs (Rheb and RhoA),1µs (Rap1A, Rap1B and Cdc42b) and 2.2µs (Rac1) of the data from analyses of equilibrium properties. The results of these analyses are discussed in the subsequent sections.

**Figure 2.**
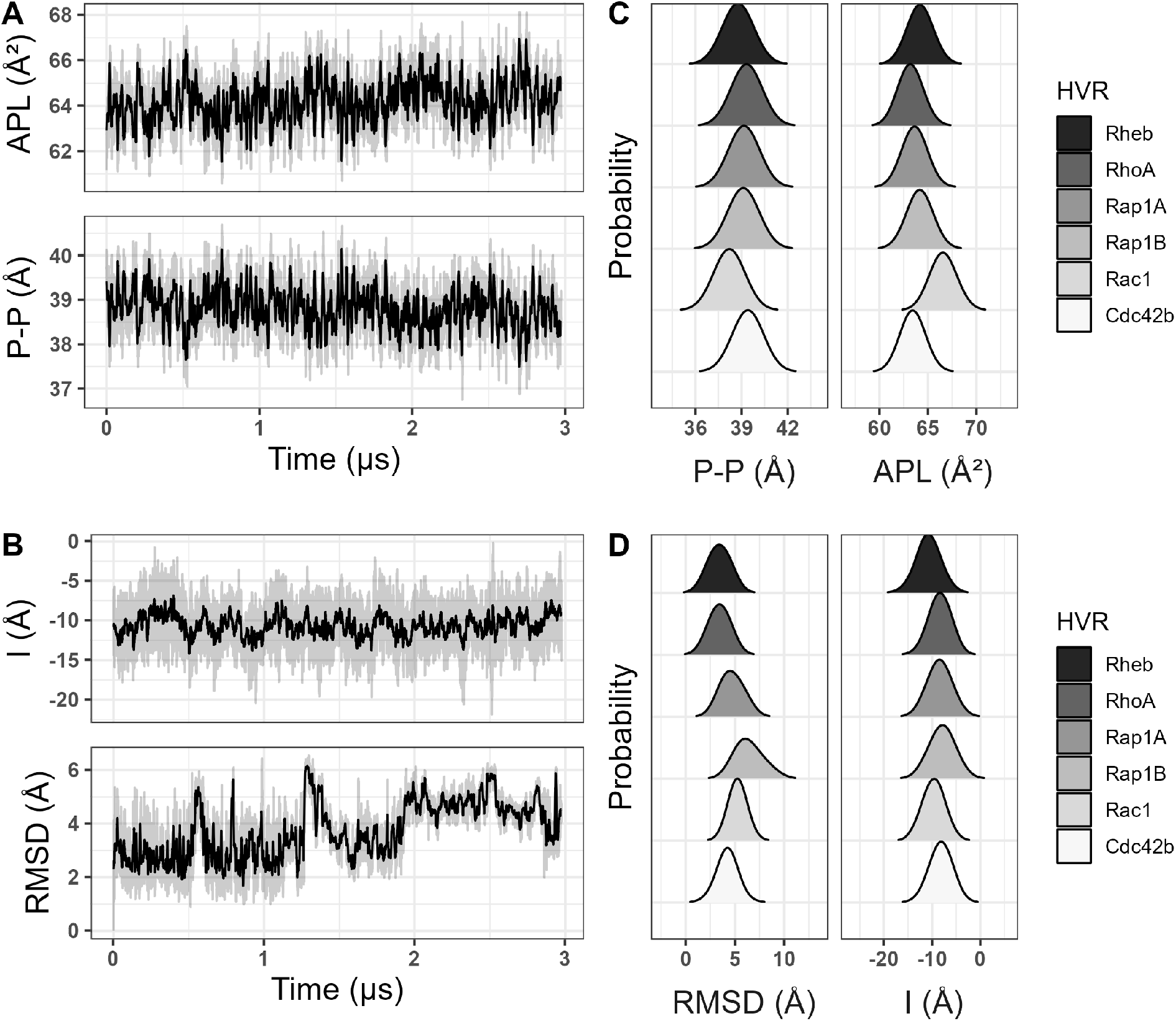
Membrane and PIDR structural properties. (A, B) Time evolution of area per lipid (APL) and bilayer thickness (P-P) in the Rheb simulation (A), and bilayer insertion depth of the prenyl chain (I) as well as backbone root mean square deviation (RMSD) for one of the Rheb peptides (B). Complete sets of plots for all systems are found in Figs S1 and S2. (C, D) Histogram of P-P and APL (C), and RMSD and I (D), from each of the six simulations. In these histograms and in all subsequent figures, data from the first 0.5µs (Rheb and RhoA), 1µs (Rap1A, Rap1B and Cdc42b) and 2.2µs (Rac 1) is excluded as an equilibration phase. For the PIDRs, averaging was done overall all three peptides except for Rac1 where only one peptide was considered (see text for details).

**Figure 3.**
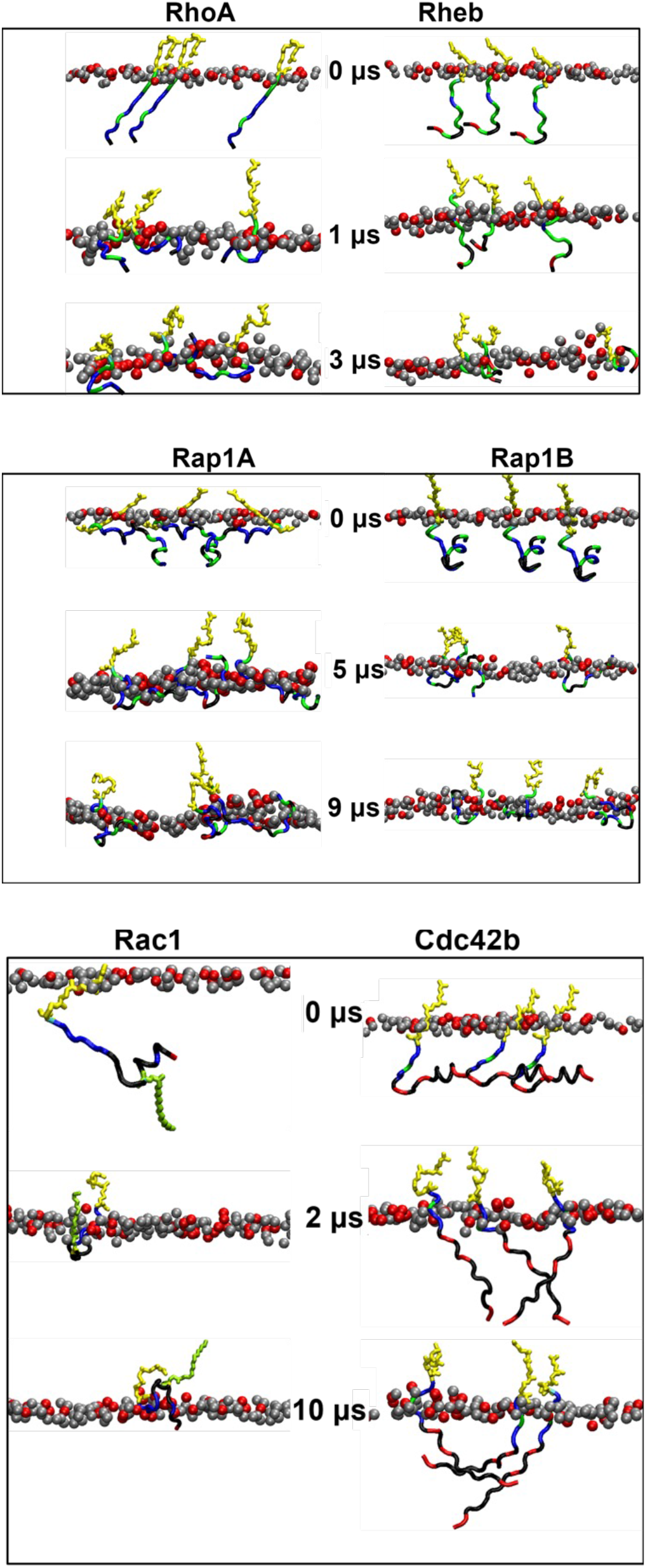
Bilayer adsorption. Shown are snapshots of the membrane-embedded PIDRs at the indicated time points from each trajectory, with phosphorus atoms of POPS in red and POPC in gray (only the peptide-hosting monolayer is shown). Prenyl chain in yellow and palmitoyl in green.

### Bilayer adsorption and membrane structure perturbation

We recently showed that PS asymmetry has only minor effects on the global structure and dynamics of a PC/PS bilayer^6^. Similar results were obtained in this work, including a slight increase in lipid packing and decrease in the lateral diffusion of lipids at the PS-containing monolayer. Since the previous analysis was based on the same lipid composition as in the current simulations, here we focus on the impact of each PIDR on bilayer structure rather than on the consequence of PS asymmetry.

The equilibrium distributions of APL and P-P displayed in Fig 2C show that there are small differences in the ensemble-averaged bilayer thickness and area per lipid among Rheb, RhoA, Rap1A, Rap1B and Cdc42 simulations. Compared with the farnesylated Rheb, the geranylgeranylatd RhoA caused a slight increase in bilayer thickness and a corresponding decrease in APL (Fig 2C). This is despite the comparable sequence length of the two PIDRs and is correlated with the deeper bilayer penetration of the longer GG tail in RhoA (Fig 2D). The two peptides also differ in bilayer adsorption, with the entire backbone of RhoA adsorbed into the headgroup region of the bilayer while part of Rheb remains in water (Fig 3). The P-P and APL distributions in the Cdc42 simulation are very similar to those of RhoA (Fig 2C) despite major differences in the length and sequence of the two PIDRs (Table 1 & Fig 1). The homologous Rap1A and Rap1B have a very similar effect on P-P or APL and only slightly differ from RhoA and Cdc42b. Moreover, there are negligible differences among the GG lipidated PIDRs in insertion depth of the prenyl chain (Fig 2D), despite notable differences in backbone adsorption (Fig 3). Specifically, the backbone of RhoA, Rap1A and Rap1B are fully adsorbed whereas all but the last few residues of Cdc42b stayed away from the bilayer (Fig 3). The dually lipidated Rac1 is unique in terms of both bilayer adsorption (Fig 3) and effect on bilayer structure, characterized by a smaller average P-P and larger APL (Fig 2C). This is due to the deep bilayer insertion of the palmitoyl chain and other hydrophobic residues in the middle of the peptide (Fig 3). Together, these results show that the bilayer structure is perturbed more by the deep penetration of the prenyl chain and, in the case of Rac1 also the palmitoyl chain, into the hydrophobic core of the bilayer. Adsorption of the PIDR backbone in the headgroup region of the host monolayer has a smaller effect on the bilayer structure.

### Structure and dynamics of PIDRs on PC-PC/PS bilayer

Fig S2 shows the overall convergence of the peptides in each system into a similar final structure. Minor divergences in some systems (e.g., RhoA and Rap1A) are not unexpected given the inherent structural diversity of PIDRs. We therefore considered all three peptides in all simulations for subsequent analysis except Rac1, where only one peptide that fully inserted into the bilayer was considered.

Consistent with their diversity in sequence (Fig 1A) and bilayer adsorption profile (Fig 3), there are significant differences among the PIDRs in the peak position and width of the time averaged *I* and RMSD distributions (Fig 2D). The broadness of the RMSD distributions also highlight significant structural diversity within each PIDR, which remained unstructured and highly dynamic during the simulations (Fig S3). For example, short helices (mostly 3_10_) were sampled only in 3% of the frames in Rap1B, 9% in Rap1A, and <2% in all others (Fig S3A). Even the PEPFOD-predicted residual helicity in some PIDRs (e.g., Rap1B and Cdc42) quickly melted during the simulations (Fig 3). Similarly, extended structures were observed only 4% of the time and only in Cdc42 (Fig S3A). The remaining simulated conformers are random coil (58-83%) or turn (11-40%) (Fig S3A). We conclude that, within the simulation timeframe, bilayer adsorption did not lead to secondary structure formation in the PIDRs studied in this work, unlike in other IDRs where such a transition has been observed^7,40–43^. However, membrane binding decreased the conformational space sampled by most PIDRs, resulting in a few well-defined conformational ensembles that are discussed later.

### PIDR bilayer localization is sequence dependent

The snapshots in Fig 3 and the location and fluctuation of individual sidechains relative to the average location of the phosphate groups in the bilayer (Fig 4) show that bilayer adsorption depends on the overall sequence composition of the PIDRs. As expected, the prenyl chain in each PIDR as well as the Palm tail in Rac1 inserted deep into the hydrophobic core, with their COM located about 10Å from the average location of the phosphate group in the peptide-bound monolayer. About five residues of each PIDR immediately upstream of the prenylated Cys reside close to the phosphate group, irrespective of sequence composition and spacing of the PBD from the site of prenylation (i.e., S). This is because all five of these residues are either basic or polar that can interact with lipids through hydrogen bonding or salt bridge formation, consistent with their localization (Fig 4). Moreover, in all cases where *S* is not zero, the intervening amino acid is Ser (Rap1A), Ser and Gly (RhoA), or two Ser residues (Rheb and Rap1B). The side chain of Ser can form a hydrogen with lipids and so can the Gly backbone, explaining their localization in the polar headgroup region. We conclude that *S* does not have a significant impact on the bilayer localization of the PIDRs studied in this work.

**Figure 4.**
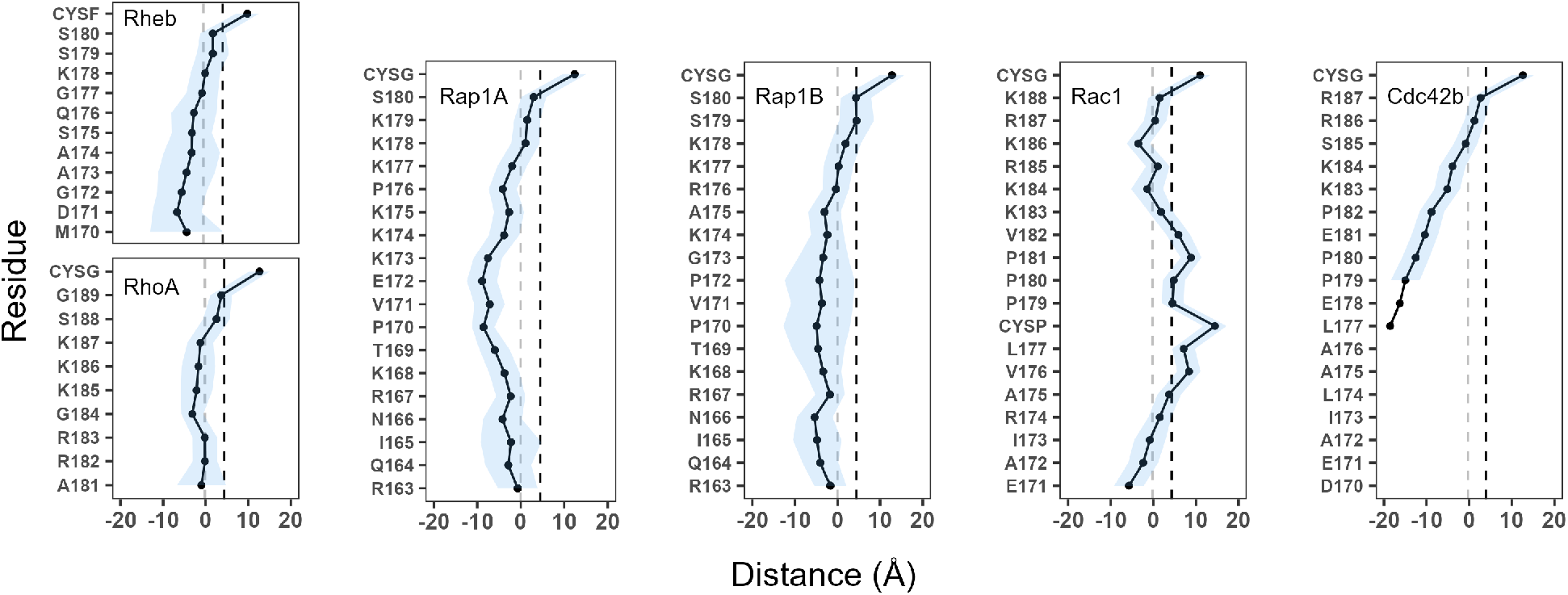
Side chain localization. Time-averaged side chain center-of-mass z-position (mean = symbols; S.D. = shades) relative to the average z-position of phosphorus atoms of the host monolayer (i.e., z-distance between sidechain COM and phosphorous atoms in the host monolayer). Vertical dashed-lines indicate the average location of the POPS phosphate (grey) and glycerol-ester oxygen atoms (black). CYSF = farnesylated cystine; CYSG = geranylgeranylated cysteine (see Table S1 for the definition of abbreviations).

Significant differences are observed in the location of the rest of the sidechains and hence the overall organization of the PIDRs on the bilayer surface (Figs 3 & 4). In the control peptide Rheb that does not have a PBD, almost all the sidechains apart from those near the prenylation site make only occasional contacts with lipids (see fluctuations in Fig 4). The lack of stable interactions is also reflected in the rather diffuse distribution of the orientation of the backbone relative to the membrane normal (Fig S4). Similarly, the PBD-containing Cdc42b engages lipids only through the C-terminal five residues, with all other sidechains staying in water (Fig 4) and the backbone lacking a specific orientation (Fig S4). This is not because Cdc42 has fewer basic residues at the PBD (Np = 4) since Rap1B with the same Np adopts a completely different bilayer organization and orientation (Fig 4; Fig S4) but rather because of the four negatively charged residues outside the PBD region (Table 1; Fig 1A). Rap1B as well as RhoA, Rap1A, and Rac1 lie nearly flat on the bilayer surface, with the sidechains at the PBD and most of those outside the PBD residing near the lipid phosphate group (Fig 4). RhoA, Rap1A and to a lesser extent Rap1B engage lipids through both C- and N-terminal residues, reflecting the presence of basic amino acids in these regions. These proteins also have orientational preferences (Fig S4). The outward bulge in the sidechain COM plot of Rap1A is caused by the single Glu in the middle of the sequence, which appears to have countered the potential impact of the neighboring non-polar sidechains (Fig 1). By contrast, Rac1’s bilayer localization is characterized by a deep insertion of the Palm tail and the neighboring hydrophobic residues, with its N-terminal Glu and other sidechains staying in water. Unlike Rap1A and Rac1 that share the same Np = 6 but differ in Nh (0.83 vs. 0.33) and B:A (3:1 vs. 1:1), Rap1B has an intermediate Nh = 0.45 and no acidic residue (B:A = 3:0). As a result, all Rap1B sidechains lie close to phosphate groups (Fig 4) and, as in RhoA, the backbone is nearly parallel to the membrane surface (Fig S4). These results show that, in addition to the PBD, the sequence composition in the rest of the PIDR play key roles in membrane engagement.

### PBD-containing PIDRs segregate lipids in a sequence dependent manner

What is the reason for, or the consequence of, the differential bilayer-organization of the PIDRs? To answer this question, we examined the overall interaction of each PIDR with PC and PS lipids. To do this, we counted the number of hydrogen bonds (N_HB_) between all polar or charged amino acids and all lipid headgroup oxygen atoms, and vdW contacts (N_C_) between all carbon atoms of non-polar residues (including prenyl and Palm chains but not proline) and acyl chains of lipids. In each case, averages were taken over three peptides (N_peptide_ = 3) in the simulation box except for Rac1 where N_peptide_ = 1. The time series of N_HB_ and N_C_ show that interactions with PS headgroup and acyl chains increased very quickly (typically within 0.1-1μs) in all the PBD-containing PIDRs (Fig S5). Interactions with PC remained unchanged or decreased over time except for Rac1 where there was an increase in N_C_ up to ∼2μs, reflecting the slow insertion of the Palm chain discussed above. Remarkably, the increase in interactions with POPS headgroups (N_HB_(PS)) is quickly followed by a corresponding increase in interactions with PS acyl chain carbons (N_C_ (PS)). This is true for all PIDRs including RhoA that does not have hydrophobic residues, but not for Rheb that lacks a PBD. This result demonstrates the key role of the PBD in segregating and clustering anionic lipids, thus generalizing previous observations on KRAS^6,27,25,32^.

As expected, N_C_ is dominated by the contribution from the lipidated cysteines and therefore its value is affected by the type of lipid modification. For example, the histograms in Fig 5 show that the average N_C_(PS) in the geranylgeranylated PIDRs varies between the narrow range of 5.8 and 7.2 but jumps to 11.2 in Rac1 because of the additional lipid modification. By contrast, N_C_(PS) = 4.1 was obtained for the farnesylated Rheb. When the lipidated cysteines were excluded, we obtained N_C_(PS) values of 0.3-0.6 for all except Cdc42b and Rac1 where it was 0.0 and 2.5, respectively. The latter values represent 0% and 22% of the total N_C_ (Fig 5B), indicating that the non-lipidated sidechains in different PIDRs contribute differently to vdW interactions. For the longer PIDRs, these contributions are loosely correlated with Nh and B:A combined but not to either separately. Moreover, the contribution of non-lipidated amino acids to N_C_ is greater in the shorter Rheb than the longer Cdc42b or Rap1A, demonstrating the importance of the overall sequence composition rather than any specific variable alone. Similarly, from the histograms in Fig 5A we find that the average N_HB_(PS) varies between 1.3 and 8.6 and N_HB_(PC) between 1.1 and 4.6. While it is clear that the PBD-containing PIDRs form more hydrogen bonding interactions with both PS and PC lipids, the extent of the interaction is not directly correlated with Np or K:R at the PBD but rather with the aggregate effect of both these variables and the sequence outside the PBD. For example, Rap1A (Np = 6, K:R = 6:0) and Rap1B (Np = 4; K:R = 3:1), as well as Rac1 (Np = 6; K:R = 4:2) and Cdc42b (Np = 4; K:R = 2:2), have comparable numbers of HB contacts with both PC and PS lipids despite their significant differences at the PBD (Fig 5A). We conclude that N_H_ and N_C_ do not correlate with just Np, Nh, K:R or B:A but rather with a complex mix of these variables.

**Figure 5.**
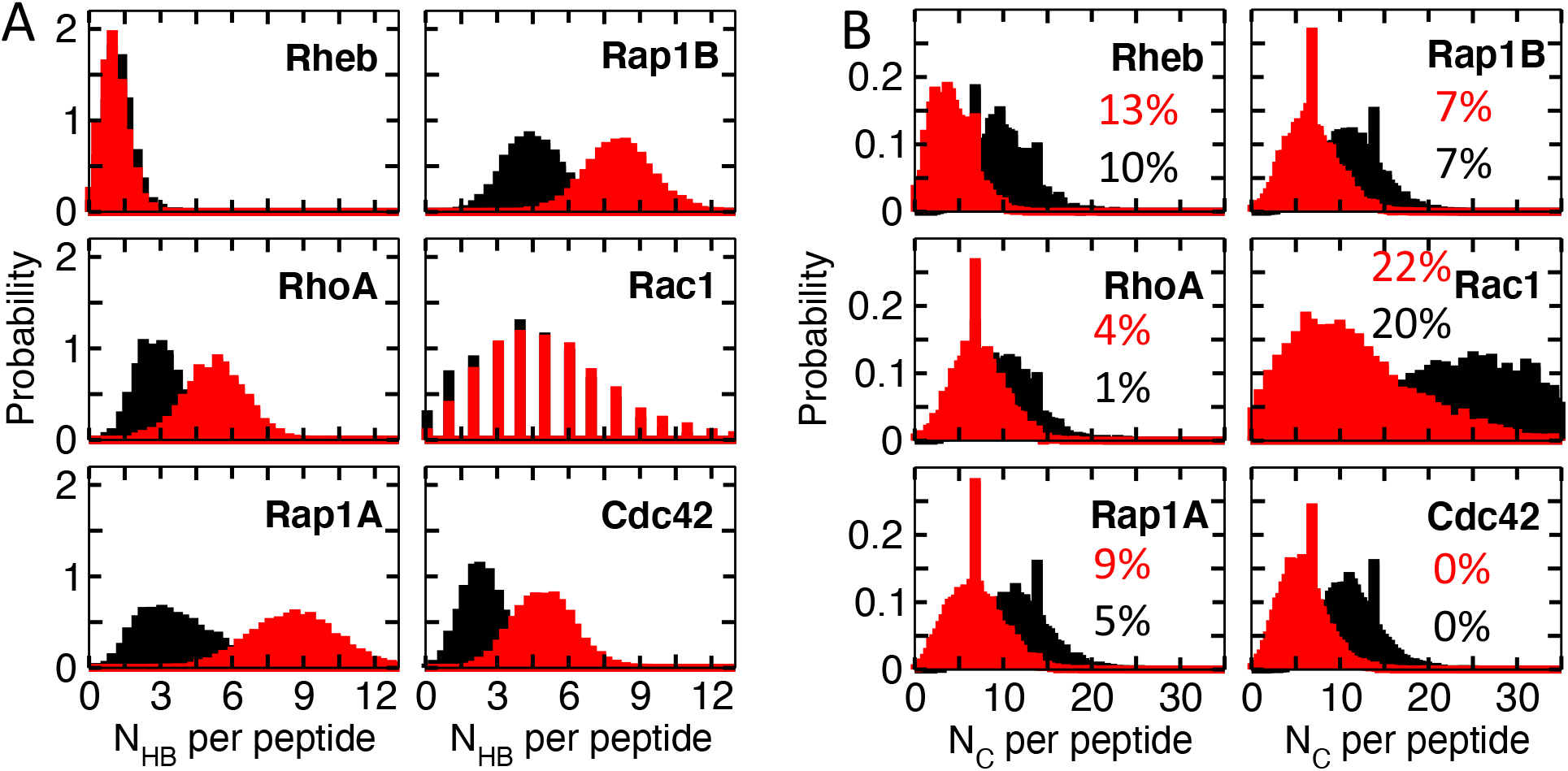
PIDR-lipid interaction. Equilibrium distributions of the numbers of hydrogen bond (N_HB_, panel A) and vdW contacts (N_C_, panel B) per peptide with POPC (black) and POPS (red) lipids. HB was calculated using a donor-acceptor distance cutoff of 3.1Å and a donor-hydrogen-acceptor angle cutoff of 30° and included all polar and charged sidechains and lipid headgroup oxygen atoms. N_C_ was computed using a carbon-carbon distance cutoff of 4Å and included all non-polar sidechain carbons (excluding proline but including lipidated cysteines) and lipid acyl chain carbon atoms. The percent contributions of non-lipidated nonpolar sidechains to N_C_ are indicated.

We previously showed that the KRAS PIDR preferentially engages anionic lipids primarily through salt bridge interactions between the basic residues of the PBD and the charged headgroup of lipids^25,32^. Similarly, Fig S6 shows a stronger interaction between Arg/Lys residues of the PIDRs studied here with the headgroup carboxyl than phosphate oxygens of POPS, except for the PBD-lacking Rheb and the deeply inserting Rac1 whose preference for phosphate and carboxyl oxygen atoms is similar. Furthermore, the equilibrium distributions of N_HB_ clearly show preference for POPS over POPC by all the PBD-containing PIDRs (Fig 5A), and this preference resulted in preferential vdW interactions (N_C_) as well. For example, for RhoA we find mean values of N_HB_(PS) = 5.3, N_HB_(PC) = 3.0, N_C_(PS) = 7.2, and N_C_(PC) = 10.1. When we consider the PC/PS ratio of 2.3 in the host monolayer, these values indicate 4-fold and ∼2-fold PS enrichment at the headgroup and acyl chain regions, respectively. Similarly, the N_HB_(PS) and N_HB_(PC) distributions for Rap1A and Rap1B suggest 5.4-fold and 4.1-fold PS enrichment at the headgroup, respectively. The PS enrichment at the acyl chain level estimated from the N_C_(PS) and N_C_(PC) distributions is ∼1.5-fold in both proteins. Even in Rac1 and Cdc42b, which represent extreme cases of PBD-containing PIDRs (see next section), PS is enriched by ∼2-fold and ∼1.2-fold at the headgroup and acyl chain levels, respectively (Fig 5). In the control Rheb, which has only one Lys and a few serine residues, the average N_HB_(PS) and N_HB_(PC) are very small and nearly equal: 1.3 and 1.1. While this suggests a 3-fold enrichment of PS at the headgroup the N_C_ distributions are consistent with the proportion of the PC/PS lipids, suggesting no lipid sorting at the acyl chain level. Notice in Fig S5 that PS lipids first engage PIDRs through HBs and then vdW contacts, and that contact with PS increases and those with PC decrease as sorting progresses. Together, these results not only demonstrate electrostatic-driven clustering and sorting of lipids at the headgroup but also show how that effect leads to a subsequent sorting of acyl chains. This finding provides the first direct evidence at the molecular level for our previous observations from EM spatial mapping and systematic lipid ablation and add back experiments^27^, where we showed that KRAS preferentially interacts with PS species with one saturated and one unsaturated acyl chain but not with di-saturated or di-unsaturated PS.

### Conformational sub-ensembles and lipid recognition

As described in *Methods*, we analyzed the simulated conformers based on a global measure (Rg or RMSD, whichever has a multimodal distribution) and a local measure based on a pseudo-dihedral angle involving four consecutive Cα atoms (Φ). For Rheb, a probability density plot of Φ versus Rg (P(Rg, Φ)) yielded at least two well separated clusters of conformers centered at P(Rg, Φ) *=* (9.29 ± 0.6Å, 111 ± 14.3°) and (10.4 ± 1.3 Å, -116 ± 27.6°) (Fig 6A). The two clusters together accounted for ∼60% of the total simulated conformers (cluster 1 = 20%; cluster 2 = 40%). The smaller size of cluster 1 is not a result of limited sampling, because each cluster was visited by all three peptides in all simulations. Conformers in both clusters of Rheb are characterized by a bent backbone conformation, with those in cluster 1 bent in the middle while those in cluster 2 twisted near the N-terminus. These geometries allowed for the polar and hydrophobic residues in the middle of the peptide (residues 173-177) to submerge in the bilayer in cluster 1 but not in cluster 2 (Fig 6B). The preponderance of more extended conformers in cluster 2 is apparent also from the distribution of sidechain locations averaged over the trajectory (Fig 4). Interactions of the single Lys with POPS is equally weak in both clusters but the location of the Gly, Ala, Ser, and Gln residues in the bilayer is conformation dependent (Fig 6B). A similar analysis of RhoA shows two dominant sub-ensembles centered at P(RMSD, Φ) = (3.61 ± 0.16Å, 45.6 ± 26.6°) and (2.44 ± 0.32Å, -87.8 ± 16.9°) populated by extended and curled conformers, respectively (Fig 6C). Each cluster accounted for 22% of the total conformers analyzed, with additional smaller clusters accounting for much of the remaining conformers. The two clusters share similarities in the bilayer insertion of the last four amino acids and in the overall interfacial localization of the remaining residues (Fig 6D). However, there are significant variations in the location and hydrogen bonding potential of many residues within the PBD, including Arg182, Arg183, Lys185, Lys186 and Lys187 (Fig 6D,E). Except for Lys186, all these residues engaged POPS better in the extended conformations (Fig 6E). This ensemble-dependent lipid recognition and variations in the PS interaction propensities of the PBD residues are consistent with previous observations on the KRAS PIDR^6,25,27^.

**Figure 6.**
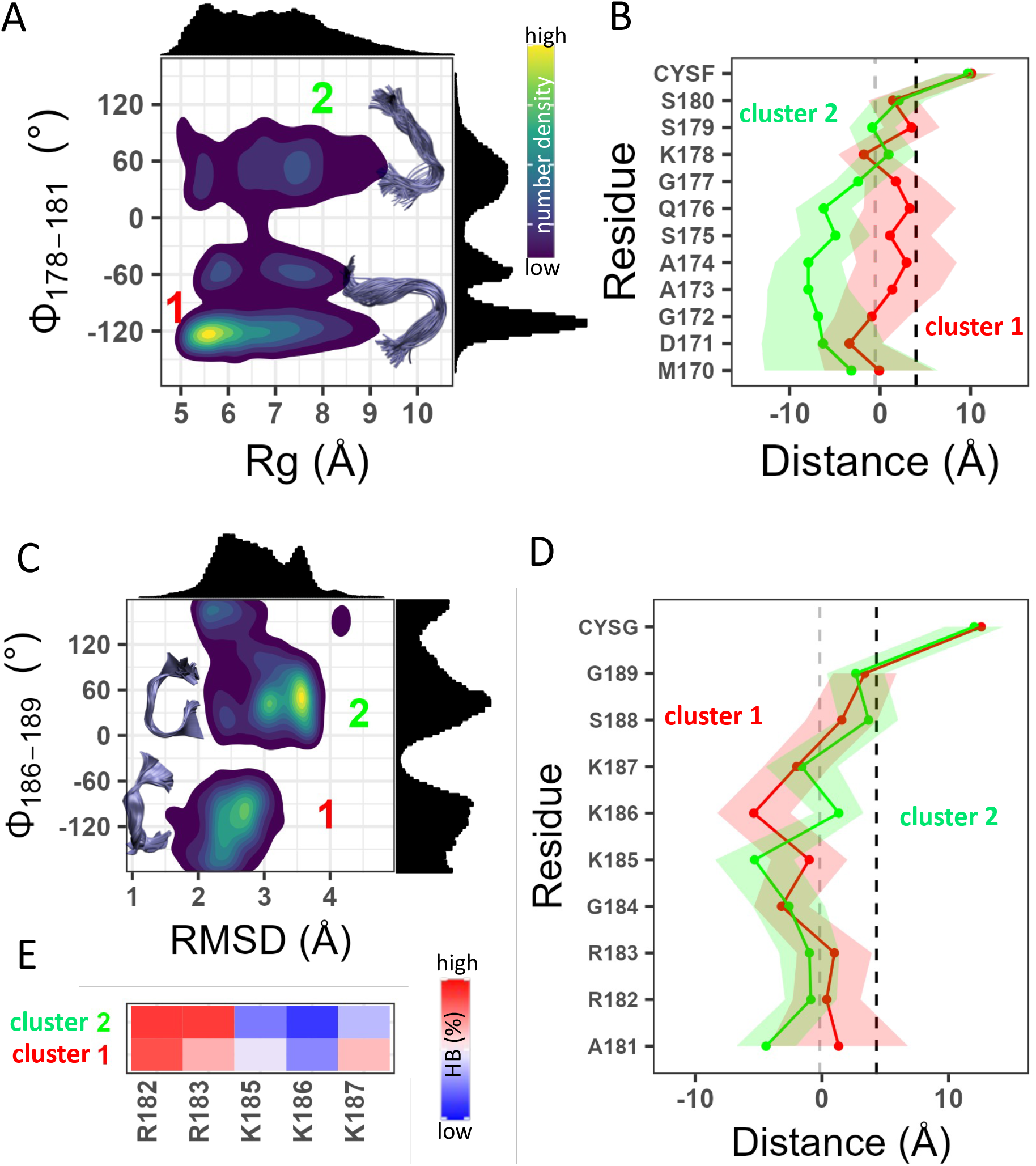
Dynamics of Rheb and RhoA PIDRs on membrane. (A, C) Normalized 2D probability density distribution of Rheb (A) and RhoA (C) based on a pseudo dihedral angle defined by the Cα atoms of indicated residues (Φ) and radius of gyration (Rg) or RMSD. Histograms of Rg/RMSD and Φ are shown at the top and right of the density plot, respectively. Insets: representative backbone conformations in clusters 1 and 2 (within 1σ of the peaks). Color scale: dark blue through yellow represent of 25% through 100% occupancy. (B, D) Time-averaged side chain center-of-mass position of each residue in cluster 1 (red) and cluster 2 (green) of Rheb (B) and RhoA (D) relative to the average phosphate z-position of the host leaflet. (E) Heatmap of normalized hydrogen bonding frequency (HBs) between the PBD sidechains of RhoA and POPS headgroup oxygen atoms. Averaging was done over ∼75,000 conformers for each system.

The homologous Rap1A and Rap1B sampled 3-4 sub-ensembles (Fig 7). In Rap1A, these included two compact (clusters 1 & 2) and two extended (3 & 4) sets of conformers (Fig 7A). The conformations in clusters 1 and 2 centered at P(Rg, Φ) = (7.8 ± 0.3Å, 122 ± 7°) and (9.6 ± 0.3Å, 122 ± 7°) differ in bilayer localization from those in clusters 3 and 4 centered at P(Rg, Φ) = (11.5 ± 0.8 Å, 49.6 ± 7.1°) and (11.5 ± 0.8Å, -96.1 ± 27°) (Fig 7B). The major difference is the deeper bilayer penetration of the N-terminal five residues in clusters 3 and 4, which includes Arg163 and Arg167. As a result, these basic residues along with Lys168 formed HBs at higher frequencies in the uncurled structures (Fig 7C). Within the PBD (residues 173-179), the frequency of HB contacts alternated between ‘hot’ and ‘cold’ in the uncurled while it is universally ‘warm’ (∼50%) in the curled ensembles (Fig 7C). Specifically, Lys173, lys175 and Lys178 barely interact with POPS in the uncurled conformers while all six lysines in the PBD partially engaged PS in the curled ones. There are also differences in HB frequency between clusters 3 and 4, which differ in Rg, and to a lesser extent between clusters 1 and 2 that differ in Φ, suggesting that lipid recognition is modulated by both global and local structural features. A similar conclusion could be drawn by comparing the two well populated clusters of Rap1B (the smaller third cluster is omitted for simplicity) (Fig 7 D-F). Centered at P(Rg, Φ) = (9.3 ± 0.6Å, 111 ± 14°) and (10.4 ± 1.3Å, -116 ± 28°), clusters 1 and 2 significantly differ in sidechain bilayer localizations in the middle of the peptide (Fig 7E) and to a lesser extent in HB frequency (Fig 7F). Specifically, Arg167 and Arg176 formed persistent HB contacts in both clusters but Arg163, Lys174, Lys177 and Lys178’s HB interactions differ between the clusters.

**Figure 7.**
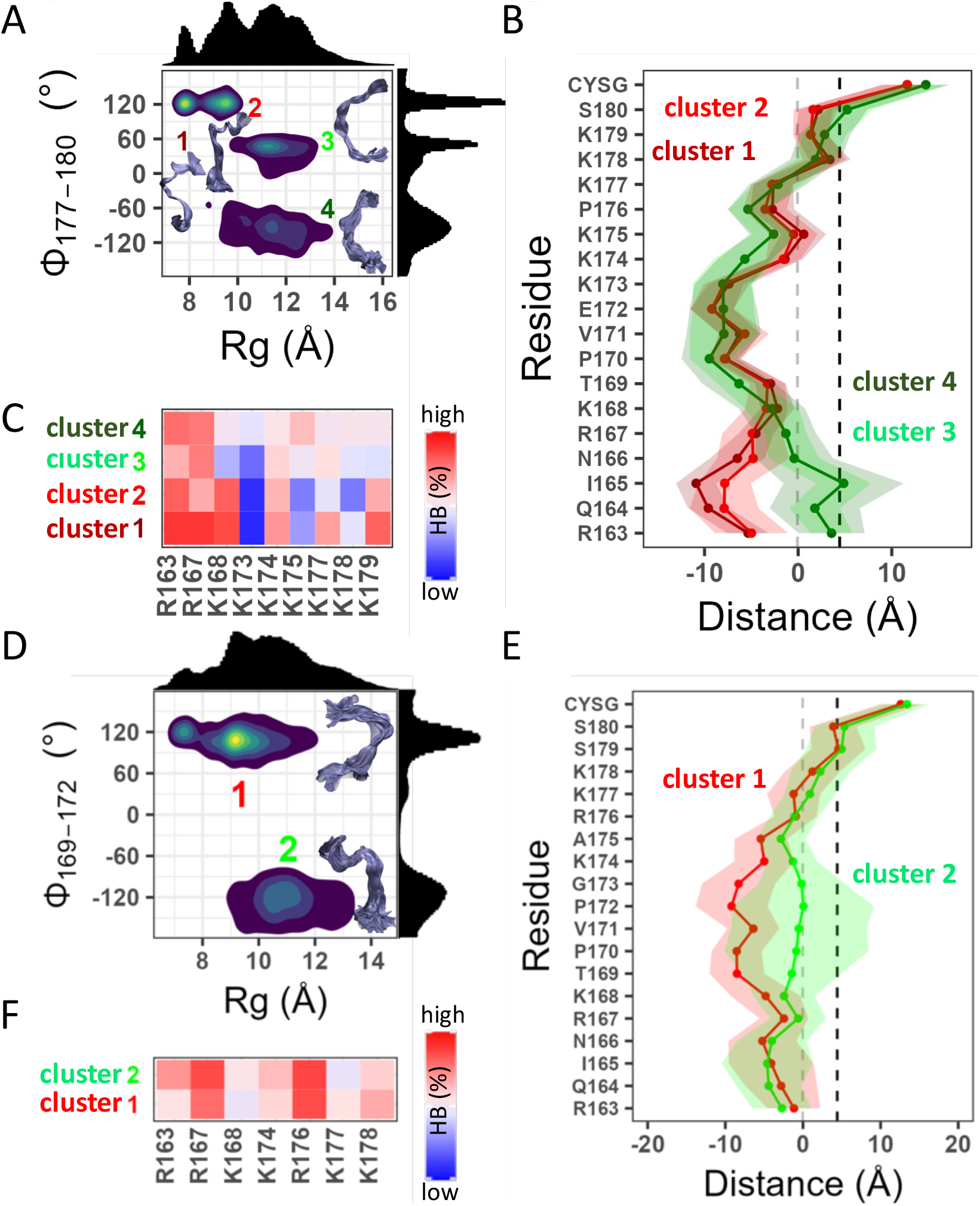
Dynamics of Rap1A and Rap1B PIDRs on membrane. Panels A-C show that the PIDR of Rap1A fluctuates between four conformational sub-states (A) that adopt two distinct bilayer adsorption profiles (B) and ensemble-dependent interactions with POPS (C). Panels D-F show that the Rap1B PIDR samples at least two distinct conformational sub-states (D) that differ in bilayer adsorption (E) and PS interaction (F). Averages were taken over ∼250,000 conformers for each protein. Color code is as in Fig 6.

The results from the RhoA, Rap1A, and Rap1B trajectories described above provide strong evidence for the correlation between backbone conformational dynamics, membrane localization, and interactions of PBD residues with anionic lipids. Our findings also highlight how the sequence composition outside the PBD modulates these correlations. A more dramatic example for the latter is provided by Rac1 and Cdc42. Despite its deep membrane penetration (Figs 3&4), the dually lipidated and more hydrophobic (Nh = 0.83) Rac1 sampled three distinct conformational sub-states (Fig 8A). Sub-states 1 and 2, which account for 36% and 7% of the total conformers analyzed, are similar in compactness but differ in backbone planarity. The less compact cluster 3 (14%) shares the same dihedral angle with cluster 2. All are characterized by a deep insertion of the center of the peptide into the hydrophobic core of the bilayer and by a stable peptide-lipid engagement (smaller fluctuations in Fig 8B). Despite these similarities, the three clusters differ in the HB frequency of several of the PBD residues (Fig 8C): Lys184 and Lys188 interact with PS more in cluster 3 than in the rest while Arg174 and Arg187 dominate the interaction in cluster 2. It is worth noting that, in all three clusters, Arg185 interacts with POPS at a higher HB frequency than the other PBD residues (Fig 8C). This may explain a previous observation where mutation of Arg185, but not the other PBD residues, to Gln resulted in a significant reduction in recruitment to the PM^26^. Moreover, the Rac1 PBD formed less frequent HBs than RhoA, Rap1A or Rap1B (Figs 5A&8C), likely because in Rac1 electrostatic interactions are complemented by a higher number of vdW contacts (Fig 5B) involving Val176, Leu177, Palm178 and Val182 that persist in all clusters.

**Figure 8.**
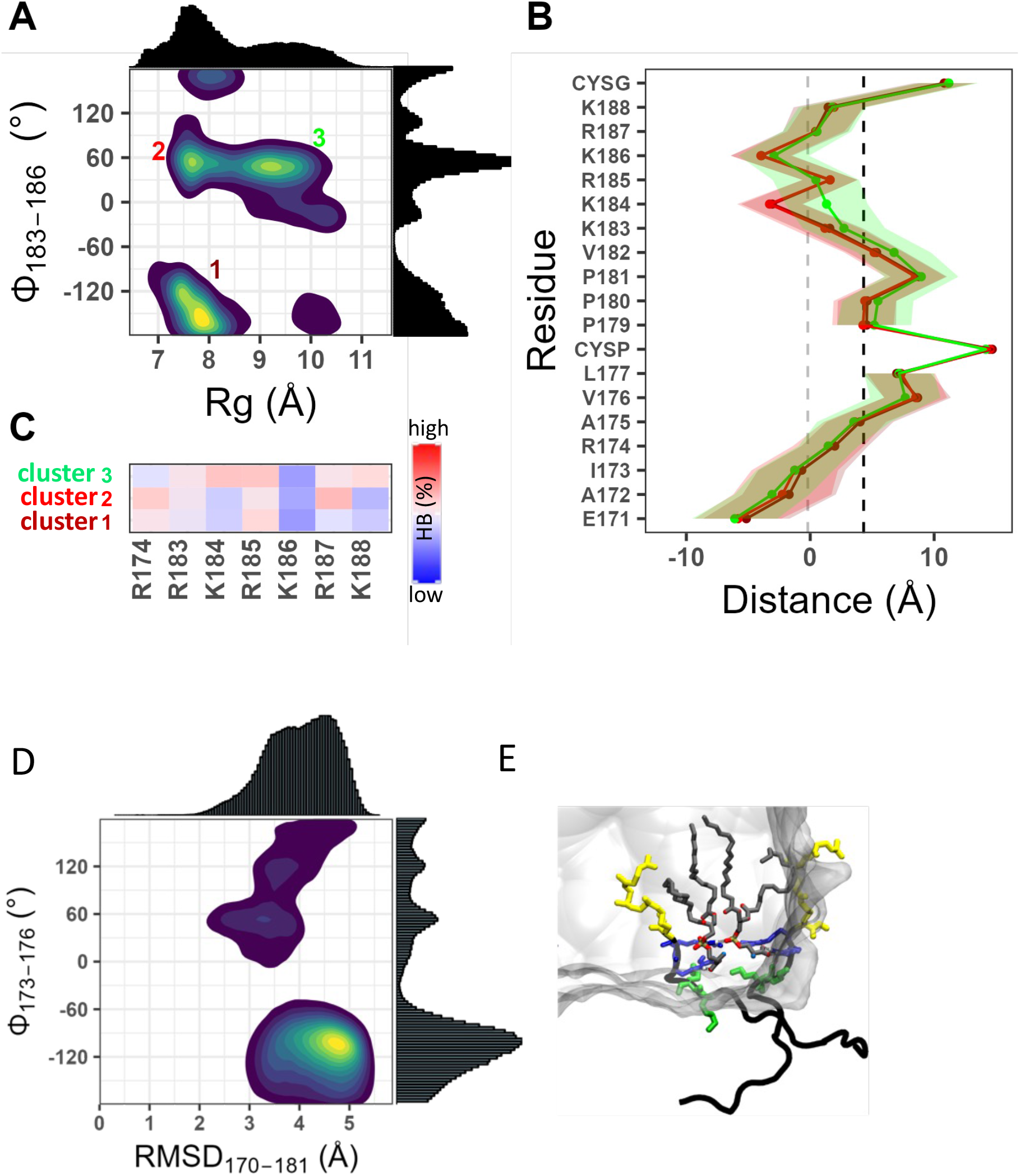
Dynamics of Rac1 and Cdc42b PIDRs on membrane. (A-C) Three clusters are observed in Rac1 (A) that differ little in bilayer adsorption (B) but differ in interaction with POPS (C). (D,E) Cdc42 does not sample defined conformational ensembles (D) with only Arg186 and Arg187 (blue) interacting with POPS while Lys183 and Lys184 (green) point away from the bilayer (E). The prenyl group is in yellow licorice and POPS in grey licorice, with oxygen atoms in red and P atoms in gold. Averaging was over ∼78,000 conformers for Rac1 and ∼250,000 conformers for Cdc42. CYSP = palmitoylated cysteine.

Cdc42b shares similarities with Rap1B at the PBD (Np = 4 in both; K:R = 2:2 vs. 3:1) but the two significantly differ in the region N-terminus of the PBD (Nh = 0.70 vs. 0.45; B:A = 0:4 vs. 3:0) (Fig 1A). Cdc42b also shares similarity with Rac1 in terms of Nh, but the latter has a Palm chain plus two additional Lys residues at the PBD while the former has three more acidic residues outside the PBD (B:A = 0:4 vs. 1:1). However, the P(RMSD, Φ) plot of Cdc42b displayed in Fig 8D shows a wide distribution of conformers that lack the well-resolved clusters observed in the other systems. Cdc42b is also unique in its bilayer adsorption and dynamics (Figs 3 & 4), and the absence of conformational preferences is reflected in the lack of specific orientational states (Fig S3). As a result of these fluctuations, only the C-terminal Arg186 and Arg187 make persistent HB contacts with POPS while Lys184 and Lys185 that often point away from the bilayer (Fig 8E) make only sporadic HB contacts. The N-terminal region remains solvated despite carrying a hydrophobic patch (residues 172-177) that is flanked by acidic residues.

## Concluding Discussion

In the current work, we used atomistic simulations to investigate six carefully selected prenylated intrinsically disordered membrane anchors of Ras and Rho family small GTPases. Ras and Rho are classical examples of molecular switches that share a conserved catalytic domain but diverge in their membrane targeting motifs and functions. The biochemical function of these proteins involves cycling of the catalytic domain between an active GTP-bound and inactive GDP-bound states. While regulated cycling between these states is required for a wide variety of physiological processes that control cell growth, motility and trafficking^44–46^, dysregulation of the cycle causes many intractable diseases including cancer^47–54^. In addition to the GTPase cycle, membrane binding is required for the cellular activity of Ras and Rho but our insight into the structure and dynamics of these proteins in membrane limited. In the past, we and others have studied the membrane insertion profile^32,38,39,55^ and energetics^56–58^ of the isolated lipid anchor (as well as the HVR^38,59^) of Ras proteins using MD simulations. We have combining MD simulations with cell signaling assays, EM, or single molecule FRET studies to show that the catalytic domain of Ras proteins interacts with membrane in multiple orientations^35,36,55,60,61^. Some of these orientations are defective in signal transduction due to occlusion of the effector-binding region by the membrane^35,62^, and reorientation is likely a general feature of prenylated small GTPases that harbor a moderately long flexible linker^8^. We have also shown that the sequence and dynamics of the KRAS lipid-anchor encode lipid selectivity^25,27^ and the conformational dynamics of the HVR is a key determinant of membrane reorientation^35,36,37^. The current work generalizes these intriguing findings by studying the RhoA, Rap1A, Rap1B, Rac1 and Cdc42b membrane targeting motifs, and thereby sets the stage for future investigations of the full-length proteins using MD, EM, and functional assays as we have done in the past^25,27,36,60,61^.

Note that RhoA, Rap1A, Rap1B, Rac1 and Cdc42b possess a polybasic domain (PBD) while Rheb does not have a PBD. In previous studies of the PBD-containing KRAS PIDR in which individual PBD residues were mutated to Gln, we observed distinct conformational distributions and patterns of interaction with anionic lipids^25,27^. Variations in conformational sampling and interaction were observed even among equally charged KRAS PBD mutants, as well as between farnesylated and geranylgeranylated KRAS anchors^27^. These findings suggested that the specific sequence composition of the PBD and the identity of the prenyl chain together determine lipid recognition by altering the conformational dynamics of the membrane anchor. The current results not only validated this hypothesis but also extended it in many respects, such as by demonstrating the significance of the amino acid sequence outside the PBD (Figs 3-8).

In RhoA, Rap1A and Rap1B, interactions of the PBD Arg and Lys residues with anionic lipids varied with the conformational ensembles of the backbone (Figs 6 & 7). In addition, not all PDB residues engaged lipids in all ensembles, with some sidechains interacting with POPS only when the backbone adopts a particular conformation. Moreover, Arg forms stronger (more frequent) salt bridge/hydrogen bonds with PS than Lys. All these observations are consistent with our previous results^25,32,27^, and demonstrate that conformation dependent interaction with anionic lipids is generalizable to prenylated PBD membrane anchors. The current results also suggest a strong correlation of interaction patterns with bilayer adsorption profiles (Fig 3), localization of sidechains (Fig 4), and orientation of the PIDR relative to the bilayer surface (Fig S4). We propose that these correlations, too, are generalizable to classical prenylated PDB anchors where the PIDR sequence outside the PBD is enriched with basic amino acids like in Rap1A and Rap1B (Fig 1A). Other examples of RAS family proteins with a classical prenylated PDB include Ral1A, Ral1B, MRAS, DIRAS1, DIRAS2, RASD1, RASl10A and RASL10B.

The Rac1 and Cdc42 membrane anchors represent two extremes of the sequence diversity in PBD-containing PIDRs. The Rac1 PIDR is enriched with non-polar residues and is palmitoylated in addition to being prenylated. While palmitoylated and prenylated PIDRs are common in small GTPases such as in NRAS, HRAS, KRAS4A, RAP2A, RAP2B and ERAS in the RAS family as well as RhoB in the Rho family, palmitoylated and prenylated PIDRs containing a PBD are rare (the only other example in the Rho family is RhoJ). Our Rac1 results suggest that conformation dependent interaction with anionic lipids also occurs in these rare PIDRs, with some variations in the details (Fig 8). We also found that unlike RhoA, Rap1A or Rap1B, membrane binding of Rac1 is dominated by vdW interactions (Figs 5 & 8). At the other end of the spectrum is Cdc42b, which has no basic residues outside the PBD but instead has four acidic residues (B:A = 0:4). For comparison, all the prenylated PBD-containing small GTPases mentioned above have a highly conserved ∼20 residues-long PIDR with a high proportion of polar plus basic (but not acidic) residues, consistent with their primary location at the anionic inner leaflet of the PM. The unique sequence composition of Cdc42b’s PIDR is responsible for its inability to fully adsorb on the PS-rich anionic model membrane (Figs 3&4) and for the lack of conformation-dependent HB contacts with lipids (Fig 8). Together, the Rac1 and Cdc42b results strongly suggest that conformation dependent lipid sorting by prenylated PBD membrane anchors is fine-tuned by the sequence composition of the flexible segment preceding the PBD. In other words, the PBD is required but may not be sufficient for a full realization of ensemble-based lipid sorting by polybasic PIDRs. This can be tested, for example, by mutating individual residues within and outside the PBD and examining their impact on membrane binding and nanoclustering using EM spatial mapping, as we have done in the past^25,27^

Additional support for the relevance of bilayer adsorption of the entire PIDR in stabilizing a set of well-defined conformational ensembles is provided by the Rheb control simulation (Fig 6). In the relatively few instances in which this peptide was able to fully adsorb on the bilayer, it adopted a more stable conformation characterized by a sharp peak of dihedral angle versus radius of gyration plot (cluster 1 in Fig 6). However, this state is sampled rarely due to the lack of stabilizing electrostatic interactions. The phase space sampled by the non-adsorbed conformers is wide (cluster 2 in Fig 6), like that of Cdc42 (Fig 8). It is possible, however, that both Cdc42 and Rheb PIDRs will sample a more defined set of conformational ensembles when bound to a bilayer of different lipid composition, in the presence of minor lipid species such as PIP2, or, in the case of Cdc42, if the acidic residues are sequestered by a binding partner. This suggests a direct relationship between the sequence of the entire PIDR and the lipid composition of the target membrane. If confirmed by additional studies such as with mutagenesis and EM spatial mapping, a major inference of significant biological relevance that follows is that lipid recognition and sorting may be encoded in the intrinsically disordered membrane anchor of *all* lipidated peripheral membrane proteins. This means that the membrane anchor is not only responsible for high affinity membrane binding but also directs the protein to the right target membrane where it participates in lipid sorting.

The current results also provide an explanation for an intriguing previous observation where we found that KRAS selectively engages PS species with an asymmetric acyl chain saturation^27^. Our finding that electrostatic-driven sorting of PS lipids at the headgroup ultimately leads to selective interactions at the acyl chain level (Figs 5&S4) provide a detailed insight into how PBD-containing membrane anchors achieve specificity for both lipid headgroup and acyl chain structures. This result has a profound implication for future studies of the cellular distribution, transport, and function of prenylated proteins with a PBD membrane anchor.

## Acknowledgements

This work was supported by the National Institutes of Health Institute of General Medicine grant R01GM144836. Computational resources have been provided by the Texas Advanced Computing Center (TACC) and Anton 2. Anton 2 computer time was provided by the Pittsburgh Supercomputing Center (PSC) through Grant R01GM116961 from the National Institutes of Health. The Anton 2 machine at PSC was generously made available by D.E. Shaw Research.

## Author contributions

A.A.G. conceived and designed the project; M.K.A. performed the simulations; M.K.A. and A.A.G. analyzed the data and wrote the paper.

## Materials & Correspondence

Correspondence and material requests should be made to AAG

