## Supplemental Table 1, Supplemental Figures 1-6 for "Conformational ensemble dependent lipid recognition and segregation by prenylated intrinsically disordered regions in small GTPases"

**Table S1: Summary of non-standard abbreviations used in the main manuscript.**

| Abbreviation | Definition | Remarks |
|---|---|---|
| PIDR | Prenylated intrinsically disordered region | Common in lipid modified small GTPases |
| PM | Plasma membrane | |
| Farn (also called SYSF) | Farnesyl | Found in Rheb |
| GG (also called SYSG) | Geranylgeranyl | Found in RhoA, Rap1A, Rap1B, Rac1, CDC42b |
| Palm (also called CYSP) | Palmitoyl | Found Rac1 |
| PBD | Polybasic domain | Found in RhoA, Rap1A, Rap1B, Rac1, CDC42b |
| Np | Number of basic residues | |
| K:R | Lys to Arg ratio | |
| Nh | The ratio of hydrophobic to total number of residues | |
| B:A | The ratio of basic to acidic residues | |
| *S* | Spacing between the last PBD residue and the prenylated Cys | |
| RMSD | Root mean square deviation | |
| Rg | Radius of gyration | |
| RC | Reaction coordinate | |
| APL | Area per lipid | Measured using the pure-POPC monolayer water box lateral dimensions. |
| P-P | Head-to-head distance | Measured as the distance between the average phosphorous atom z-position in the two leaflets of a bilayer. |
| HB | Hydrogen bond | |
| $N_{HB}$ | Number of hydrogen bonds | |
| $N_C$ | Number of hydrophobic (carbon-carbon) contacts | |

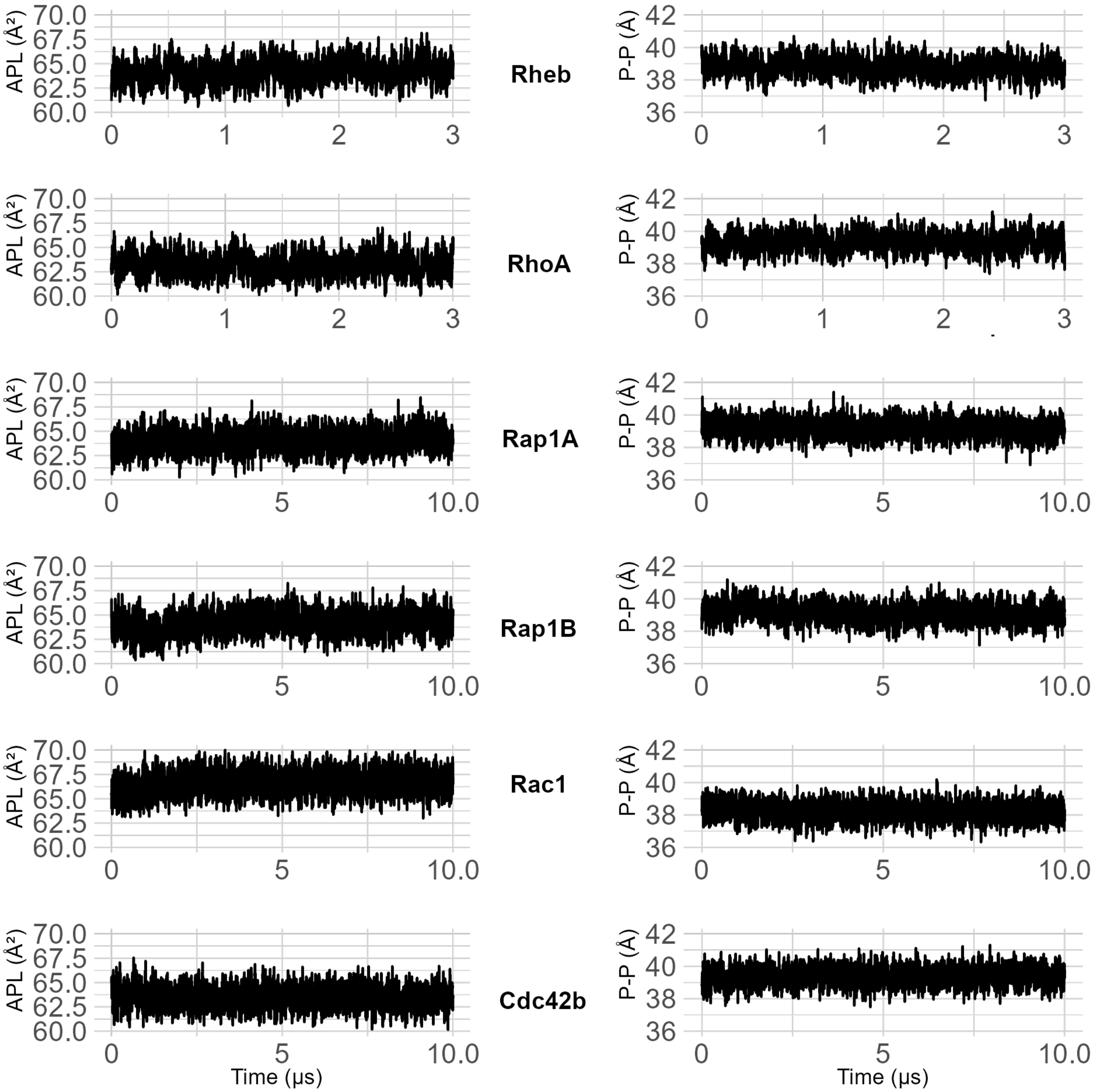


**Figure S1.** Time evolution of area per lipid (APL) and bilayer thickness (P-P) for each of the Rheb, RhoA, Rap1A, Rap1B, Rac1 and Cdc42b PIDR simulations. P-P was measured as the average distance along the membrane normal between phosphorous atoms at two the leaflets. The Rheb and RhoA simulations were 3µs long and the rest 10µs.

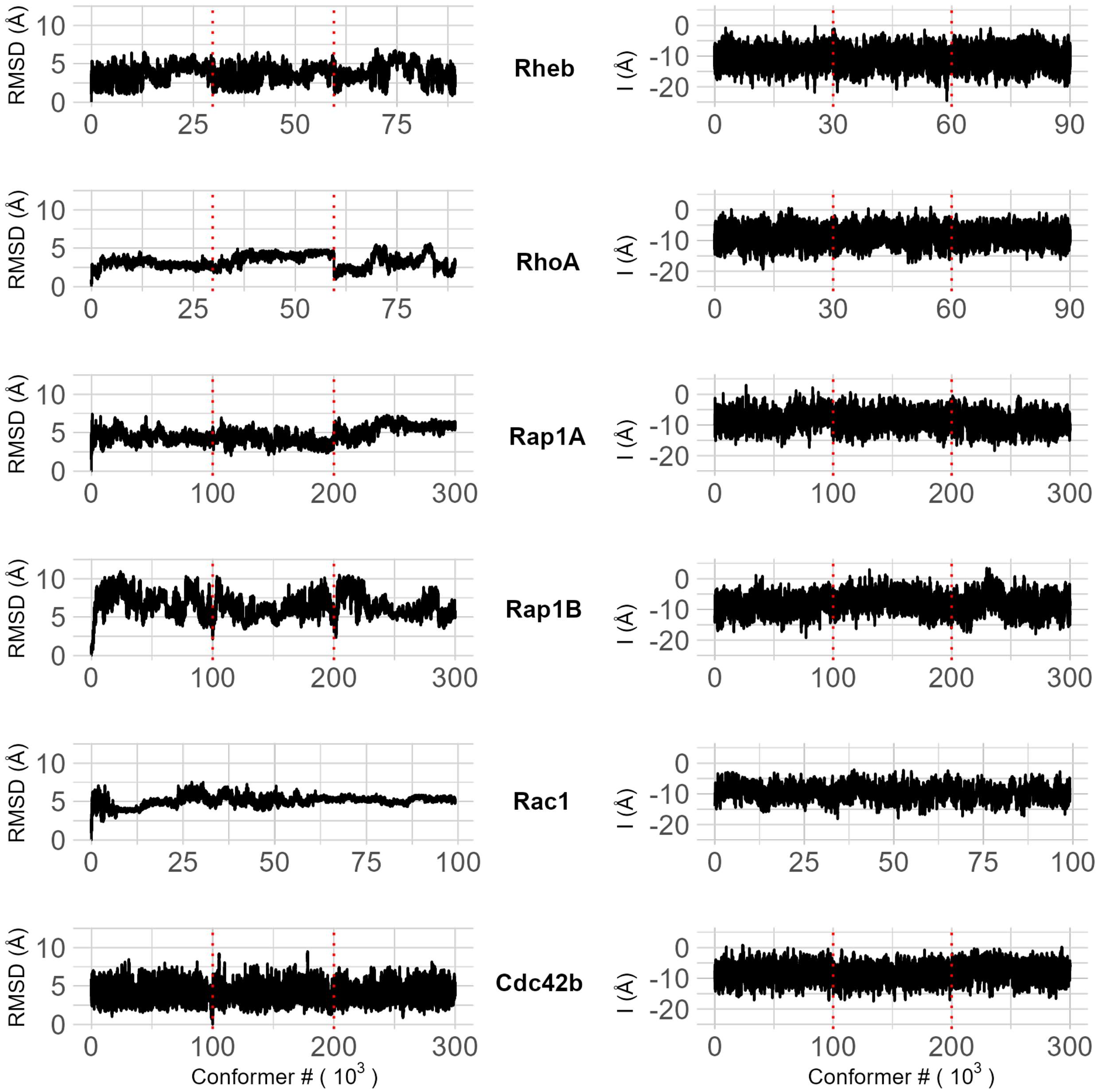


**Figure S2.** Root-mean-square deviation (RMSD; left) and membrane insertion depth (I; right) of of the simulated PIDR peptides. RMSD was measured using the initial structures as reference, and I was measured using the center-of-mass of the prenyl chain (z-distance of the center-of-mass of the prenyl chain from the bilayer center). Vertical dotted lines demarcate the three peptides in each simulation except Rac1 where only one peptide was considered (see main text for explanation). Trajectories were sampled every 100ps.

A

| Structural element | Rheb | RhoA | Rap1A | Rap1B | Rac1 | Cdc42b |
|---|---|---|---|---|---|---|
| β-bridge | 0.2 | 0.0 | 0.1 | 0.4 | 1.0 | 0.8 |
| Extended | 0.0 | 0.0 | 0.0 | 0.0 | 0.0 | 3.7 |
| Coil | 58 | 78 | 60 | 63 | 62 | 83 |
| Turn | 40 | 21 | 31 | 33 | 37 | 11 |
| $3_{10}$-helix | 1.6 | 0.1 | 6.1 | 2.0 | 0.0 | 0.4 |
| α-helix | 0.0 | 0.0 | 2.9 | 1.0 | 0.0 | 0.8 |

B

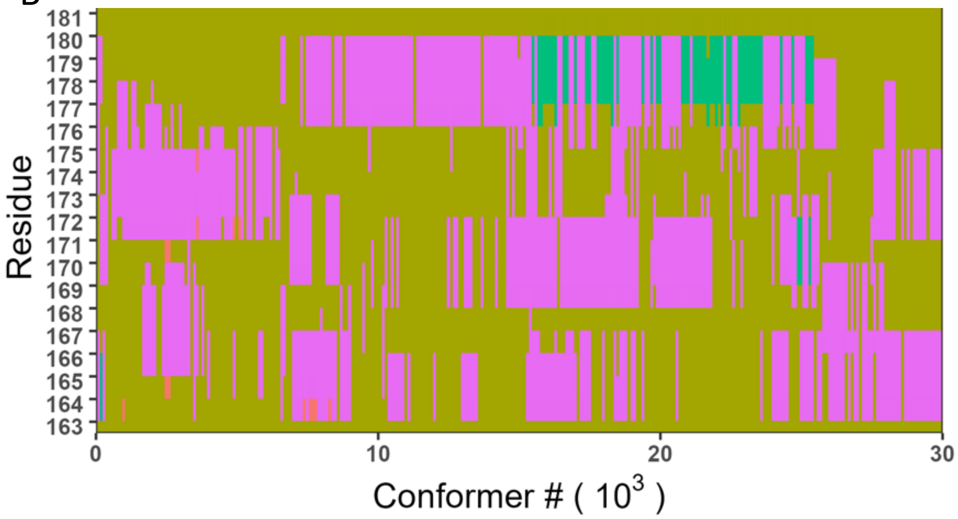


**Figure S3**. The PIDRs remained unstructured during the simulations. (**A**) Percentage of backbone secondary structure elements sampled during the simulation of each PIDR, showing that the simulated conformers are predominantly unstructured (random coil and turn). (**B**) An example of the time evolution of secondary structure elements (using Rap1A; color code as in panel **A**), showing that the peptide fluctuates between random coil and turn with rare transitions to helical conformations (sampling frequency was 1ns and concatenated data for the three peptides was used).

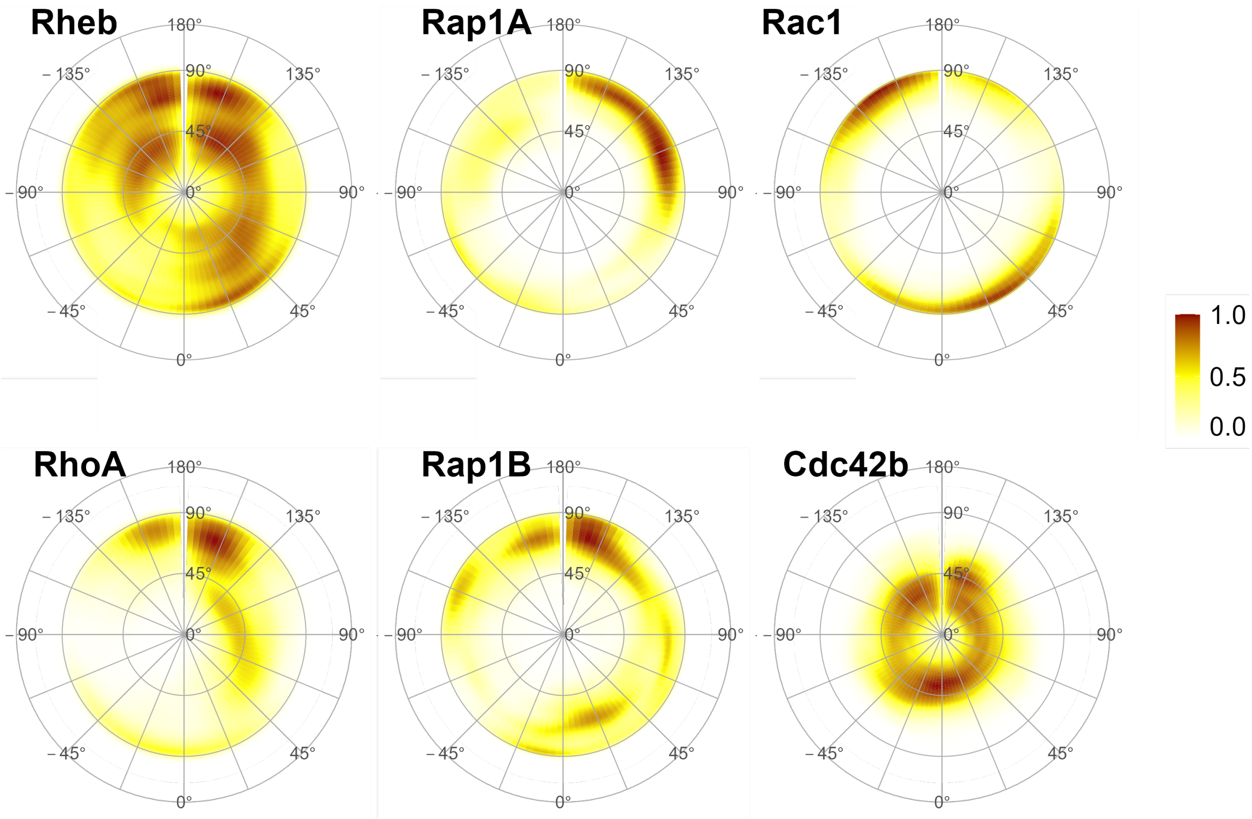


**Figure S4**. Normalized density maps of backbone tilt and rotation angles with the tilt angle [0, 90] shown along the radial coordinate and rotation and along the angular coordinate [−180, 180], quantifying the orientation of each PIDR relative to the bilayer normal.

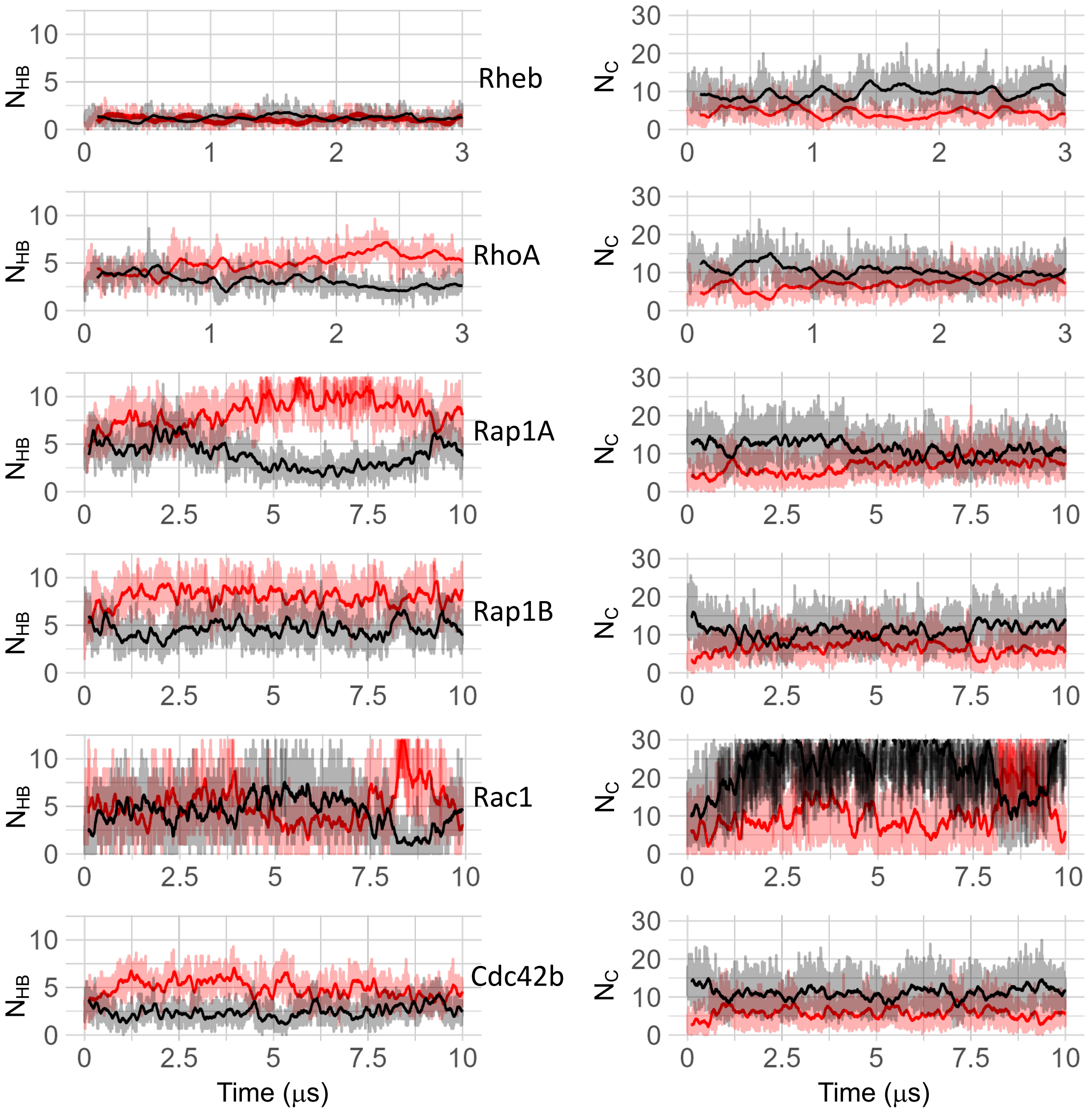


**Figure S5.** Time series of numbers of hydrogen bonds per peptide ($N_{HB}$, left panels) and vdW contacts per peptide ($N_C$, right panels) with PS (red) and PC (black) lipids. HB was calculated using all charged and polar sidechains and lipid headgroup oxygen atoms, and $N_C$ using carbon atoms of non-polar sidechains (excluding proline) and lipid acyl chains. Data sampled every 1ns is shown in lighter shades and 100ns running averages in darker shades.

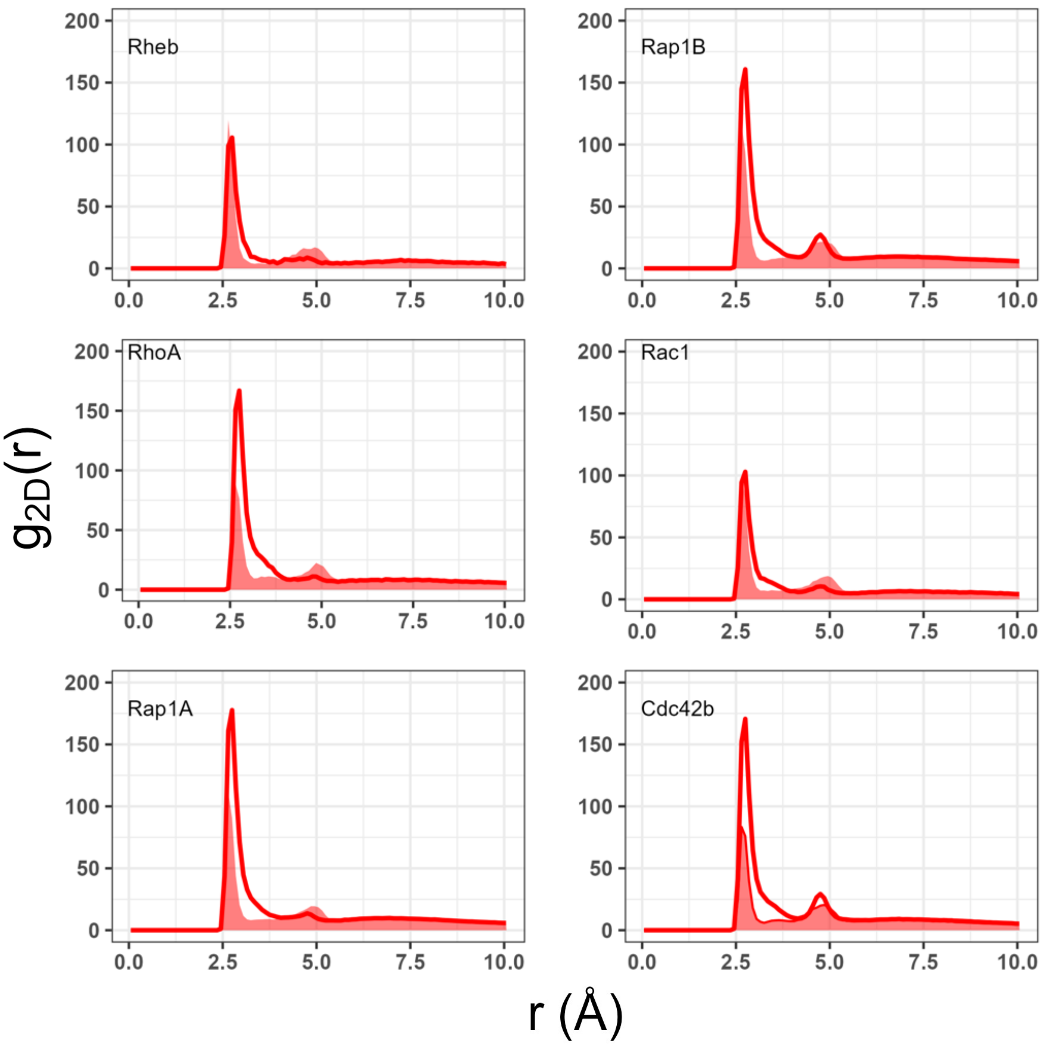


**Figure S6.** Two-dimensional radial pair distribution of POPS carboxyl (lines) and phosphate (shaded) oxygen atoms around Lys and Arg side chain nitrogen atoms of each PIDR. Both oxygen atoms of the serine headgroup (O13A and O13B in Charmm), the two phosphate oxygen atoms that are not bonded with carbon (O13 and O14), one of the terminal nitrogen atoms of Arg (NH1), and Nz of Lys were used for the calculations.
